# kakapo: Easy extraction and annotation of genes from raw RNA-seq reads

**DOI:** 10.1101/2023.02.13.528395

**Authors:** Karolis Ramanauskas, Boris Igić

## Abstract

`kakapo` (kākāpō) is a python-based pipeline that allows users to extract and assemble one or more specified genes or gene families. It flexibly uses original RNA-seq read or GenBank SRA accession inputs without performing assembly of entire transcriptomes. The pipeline identifies open reading frames in the assembled gene transcripts and annotates them. It optionally filters raw reads for ribosomal, plastid, and mitochondrial reads, or reads belonging to non-target organisms (e.g., viral, bacterial, human). `kakapo` can be employed to extract arbitrary loci, such as those commonly used for phylogenetic inference in systematics or candidate genes and gene families in phylogenomic and metagenomic studies. We provide example applications and discuss how its use can offset the declining value of the GenBank’s single-gene databases and help assemble datasets for a variety of phylogenetic analyses.

## Introduction

A variety of biological problems necessitate analyses of genetic sequence data in both basic and applied research. Until recently, there were two primary sources of low-throughput sequence data: newly generated sequences in a laboratory and sequences deposited in public databases, such as GenBank. In the past 20 years, a qualitative transition has taken place, driven by a technological shift in the kind and amount of data generated. The Sequence Read Archive (SRA) for high-throughput, massively parallel, sequencing data at the National Center for Biotechnology Information (NCBI) was established in 2009 (Kodama et al. 2012). The size of the SRA, administered by the National Center for Biotechnology Information (NCBI), has since snowballed. It has approximately doubled (to 20 petabytes) just in the last three years, between 2019 and 2022 (Katz et al. 2022). In the meantime, the relative value of single-gene submissions to GenBank’s non-redundant (nr) database has declined, despite its continued growth, as reliance on the inexpensive and large datasets produced by the new sequencing tools takes hold in many areas of biology.

Currently, about one-third of the SRA database is comprised of RNA-seq data, representing a wide range of taxa and genes appropriate for a variety of analyses at the intersection of genomics and evolution, such as those broadly referred to as “phylogenomics” and “metagenomics” (Eisen et al. 1997; Handelsman et al. 1998). While such data is free and publicly available, it is not necessarily easily accessible. For example, on NCBI’s website, only a subset of all SRA datasets can be queried with a sequence-search-tool BLAST, and assemblies are not required to be uploaded along with raw reads. Because of these and other hurdles, the weight of rapidly changing and tedious tasks required to process raw read data, often falls on individual researchers. In our experience, many groups are not equipped to take on the significant required diversion to the research program to both enable the use of new tools for sequence analysis and their constant updating. Therefore, the complexity of the analyses and the lack of integrated tools to conduct them comprise key impediments for the implementation of phylogenomic methods.

kakapo is a modular pipeline intended to remove the barriers to entry for many kinds of phylogenomic analyses. It relies on established tools and stitches them together in a reproducible, self-documenting fashion. The flow of the pipeline is fully defined using one or more configuration files, which are provided as templates with reasonable default values. Without additional intervention, it stores intermediate results and does not repeat previously completed analyses. We have already employed kakapo in published work (Sánchez-Cabrera et al. 2021; Ramanauskas and Igić 2021). Here, we describe the implementation of key steps in the pipeline, and provide readily reproducible examples, which showcase its modular features.

## Implementation

kakapo is open source software, released under the GNU General Public License v3.0 and available at https://github.com/karolisr/kakapo. The pipeline requires Python version 3.8 or higher. It can be installed or upgraded on machines running GNU/Linux or macOS operating systems, using the Python package installer pip) by running the following command in a terminal window:

~~~
pip install --user --upgrade git+https://github.com/karolisr/kakapo
~~~

Once installed, the program is executed by typing `kakapo`, which prints a brief usage reference. kakapo checks if the user’s system contains the required dependencies. If any of the dependencies are not found on the system, it attempts to install them automatically to the standard application data directory `$HOME/.local/share/kakapo`. This is done by running:

~~~
kakapo —install-deps
~~~

To increase repeatability and reproducibility, a run can be performed with `--force-deps` option, which ignores any dependencies found on the user’s system. This ensures that the same software versions are used for analyses performed at different times, without regard to the presence of the different versions that may be installed systemwide.

### Process RNA-seq reads

The pipeline minimally requires the project configuration file.

~~~
kakapo --cfg project_configuration_file
~~~

The provided template contains broadly appropriate default values and arguments, so that the user should only need to provide any combination of three types of “targets”: (1) SRA accessions, (2) paths to FASTQ files (plain or compressed), (3) a list of FASTA files containing previously assembled transcriptomes (see “Targeted transcript assembly” below). This step then downloads the FASTQ files and the associated metadata for the SRA accessions. Next, both types of FASTQ files— the ones provided by the user and the ones downloaded from NCBI—are processed identically as follows. First, the reads are optionally processed using Rcorrector (Song and Florea 2015) to correct for random sequencing errors. The reads are then trimmed using Trimmomatic (Bolger et al. 2014) to remove bases with low Phred scores, low quality reads, and any adapter or other Illumina-specific sequences.

### Filter RNA-seq reads

Often, RNA-seq reads contain RNAs from unintended or unwanted subjects, such as non-target organelles, microbial infections, and human contaminants. Therefore kakapo provides optional filtering of reads in two different ways.

#### Filter with Bowtie 2

In the first step kakapo uses Bowtie 2 (Langmead and Salzberg 2012) to optionally map the reads to reference genomes. Based on the taxonomic classification, kakapo automatically determines which DNA-containing organelles may be found in the sample and constructs a database from the appropriate RefSeq genomes by gradually broadening the taxonomic scope until at least one genome is found. If more than ten appropriate RefSeq genomes exist, a subset of ten of them is selected randomly. Alternatively, if the default behavior is undesirable, the user can override it by providing FASTA files of plastid and/or mitochondrial genomes. Any number of additional reference sequences can be provided as well.

In addition to the Bowtie 2 output in SAM format, kakapo produces a separate set of FASTQ files for each given reference. This is useful if, for example, the sample is suspected to be infected with—and contains reads from—a known pathogen.

#### Filter with Kraken 2

The second filtering step uses Kraken 2 (Wood et al. 2019), and performs rapid taxonomic classification of short reads using prebuilt reference databases. By default, kakapo downloads six prebuilt Kraken 2 databases. For Small and Large Subunit ribosomal RNA filtering, kakapo uses 16S_Silva138 database made available by the Kraken 2 developers at https://ccb.jhu.edu/software/kraken2. This database is constructed from a diverse set of rRNA sequences provided by SILVA (Quast et al. 2013). For bacterial, archaeal, viral, and human read filtering, Kraken 2 developers provide `minikraken_8GB` database. Additionally, four custom-built databases, derived from RefSeq libraries, are provided: `mitochondrion_and_plastid, mitochondrion, plastid, and viral`. The user can optionally create custom Kraken 2 databases, to filter plasmid, protozoan, fungal, and other reads.

### Filtered Read Set: Alternative Stopping Point

Using only the steps outlined above, the user can use the generated output (a filtered set of FASTQ files) as a largely contaminant-free input dataset for downstream transcriptome assembly. If so, at this point in the workflow, the output can be found in a directory named using the SRA accession or a FASTQ file name provided in the configuration file:

~~~
[OUTPUT_DIRECTORY]/01-global/05-kraken2-filtered-fq-data/[SAMPLE_NAME]
~~~

For convenience of those users who simply want to use kakapo to process raw RNA-seq reads before using the assembler of their choice to perform full transcriptome assemblies, a `--stop-after-filter` option can be provided, which terminates the pipeline at this point.

### Produce BLAST databases for filtered RNA-seq reads

kakapo performs two additional steps on the filtered RNA-seq reads. First, seqtk (Li 2012) is used to convert the FASTQ files to FASTA files. Second, BLAST (Altschul et al. 1990) databases are generated using `makeblastdb` (included with BLAST+ set of executables) from the FASTA files generated above.

### Process query sequences

kakapo uses a second type of configuration file in which the user can define one or more “search strategy” entries for the genes they wish to search for in the RNA-Seq reads. Each search strategy entry is intended to encapsulate the information about a gene or a gene family, which can then be used by kakapo to find matching RNA-seq reads and assemble transcripts of interest in a targeted manner. The user can tune the parameters of a search strategy to make the search as narrow or as wide as needed. Each search strategy entry can contain a combination of any number of Pfam (Bateman et al. 2002) family and NCBI protein accessions, NCBI Entrez query, and a set of user provided FASTA files. kakapo then combines the amino acid sequences obtained from different sources, and prunes the set based on the sequence identity and length range limits set by the user (also defined within a search strategy entry).

In order for kakapo to process query sequences and to perform the subsequent search, the user needs to provide the path of the search strategies file in addition to the project configuration file described previously:

~~~
kakapo --cfg project_configuration_file --ss search_strategies_file
~~~

### Search for RNA-seq reads

To find the candidate RNA-seq reads matching the user provided query, kakapo first runs `tblastn` (Altschul et al. 1990) using the BLAST databases that were built earlier in the pipeline. The parameters used by `tblastn` can be changed in the project configuration file. However, the settings provided in the template file have been observed to work well in all the cases we tested. When tuning BLAST parameters for this step, the user should err on more permissive values; the parameter values that produce more hits ought to generally be considered superior. This includes larger thresholds for `evalue, max_hsps`, and `max_target_seqs` parameters, and lower `qcov_hsp_perc` threshold.

To enrich the set of potentially matching reads, the set of hits from the previous step is passed to VSEARCH (Rognes et al. 2016) and used to search the original FASTQ files. Because separate queries may match the same read, the resulting set of reads is checked for duplicate ids and only the reads with unique ids are retained.

The resulting set of reads is a targeted subset of the original sample, biased for reads with higher than average sequence similarity to the query sequence.

### Targeted transcript assembly

To assemble the transcripts, kakapo runs rnaSPAdes (Rognes et al. 2016) using the reads from the previous step as input. The resulting set of transcripts (if any) may or may not be a good match for the gene of interest, to find only those transcripts that match the provided query sequences, `tblastn` is used with the parameter values specified in the search strategies file. If a list of FASTA files containing previously assembled transcriptomes was provided by the user, they are also searched at this step.

### Transcript annotation

Next, kakapo determines the genetic code for each sample, based on the sample origin (NCBI TaxID) and the genomic source (nucleus, chloroplast (if applicable), mitochondrion). Specifically, the transcripts originating from the reads that mapped to plastid or mitochondrial genomes are assigned the correct genetic code. This is followed by a search for open reading frames (ORFs) in the assembled transcripts, regardless of whether transcriptome assembly was performed using kakapo or provided by the user. Alternative ORFs are classified based on parameter settings in the relevant search strategy entry. The highest scoring ORF is labeled as a coding sequence (CDS). CDS is translated and can be optionally piped through InterProScan 5 (Jones et al. 2014) for functional annotation.

At the end, kakapo produces a series of FASTA files (with associated annotation files in GFF format, where appropriate), containing:

- complete transcript sequences (nucleotide FASTA)
- annotations for the transcripts (GFF)
- extracted ORFs (nucleotide FASTA)
- extracted ORFs (amino acid FASTA)

## Example Data and Analyses

### Data

The dataset consists of previously generated reads from a single individual of common house cactus, *Schlumbergera truncata* (“the false Christmas cactus” or “crab cactus”; Ramanauskas and Igić 2021). The species is commonly sold and traded, especially near the winter solstice. It is a short-day flowering, clonally reproduced epiphyte native to Brazil. It is also critically endangered in its native habitat (Taylor and Zappi 2017).

We extracted RNA from a single style—approximately the size of 1/3 of a toothpick—sampled from a *S. truncata* individual accession 19SF1, obtained from the Chicago botanist Joey Santore, of *Crime Pays but Botany Doesn’t.* Sequencing library was prepared using the KAPA Stranded mRNA-Seq (Roche). The sample was one of eighty-seven total samples sequenced on a single lane of Illumina NovaSeq 6000 platform (paired-end 150 bp reads) at the Duke University Center for Genomic and Computational Biology.

Both example uses employ the same reads, generated from this sample. First, we demonstrate a phylotranscriptomic use, in which we employ a candidate-based approach to sequence and characterize genes responsible for self-incompatibility. Second, we demonstrate metatranscriptomic use, in which we assemble a genome of a common plant pathogen that afflicts the same individual.

kakapo distribution includes an archive of the example dataset with configuration, input, and output files.

### Finding candidate genes in the RNA-seq haystack: Self-incompatibility RNases

Flowers of many plants, including many cacti, are hermaphrodites—they contain both male and female function. Therefore, individuals can potentially self-fertilize, but most do not. They instead express a genetic mechanism that sorts incoming pollen, rejecting all pollen that matches a specific genomic region. One widespread self-incompatibility mechanism partly relies on ribonucleases (RNases), specifically those from the T2/S-RNase protein family, termed “S-RNases.” In many conservation projects, it is helpful to know the genotype at this locus of individuals that are about to be transplanted to prevent a situation in which reduction of diversity among S-RNases causes cross-sterility and reproductive failure.

### Determining the identity of a plant pathogen: Cactus Virus X

The stems of *Schlumbergera truncata* individual, subject of our study, were unusually purple. As is the case with other commercially grown and clonally reproduced plants, individuals may acquire a variety of non-lethal chronic infections. Many pathogens may cause similar symptoms, or the infections remain asymptomatic. We use kakapo to interrogate the possible causes of a viral infection in Joey Santore’s crab cactus. One possible culprit is a well-known ssRNA virus group, broadly termed “Cactus Virus X” (Alphaflexiviridae). Alphaflexiviruses have a single-stranded, positive-sense RNA genome, variable in length but generally approximately 5-9kb (Kreuze et al. 2020).

### kakapo *configuration*

kakapo was configured to run with all the optional read processing steps turned on: Rcorrector, Bowtie 2, and Kraken 2. Bowtie 2 was set to filter mitochondrial and chloroplast reads. To check for the presence of Cactus virus X reads, the complete genome of Cactus virus X (GenBank accession NC_002815.2) was used as an additional reference for Bowtie 2 filtering step. To search for S-RNases, we prepared search strategies for T2/S-RNase gene family members. Additional search strategy for a control gene Elongation factor 1-*α* 1 or eEF1a1 was prepared. eEF1a1 was highly expressed in all tissues tested in Ramanauskas and Igić (2021). To determine and compare the expression levels of KAKAPo-recovered genes, RNA-seq reads and assembly from Ramanauskas and Igić (2021) was used, which included nine *S. truncata* individuals (15H01–15H09).

## Results

### Genotyping Schlumbergera truncata 19SF1 at the Self-incompatibility Locus

Using the T2/S-RNase search strategy kakapo produced eight distinct transcripts. Of these, four contained coding sequences near expected length. Each of the four sequences were identical across assembled lengths to *Schlumbergera truncata* RNases in other individuals, previously described in Ramanauskas and Igić (2021). Two of these matched S-RNase alleles S1 and S2, which were previously sequenced. The remaining two matched non-S-locus (Class I and Class II) RNases, members of this large protein family (Figure 2a). In Figure 2b, we show that the expression of these two transcripts is high, we fail to detect other alleles, and implication is that the plant ought not be crossable with 15H01, 15H02, or 15H04, but ought to cross readily with other genotypes.

**Figure 1:**
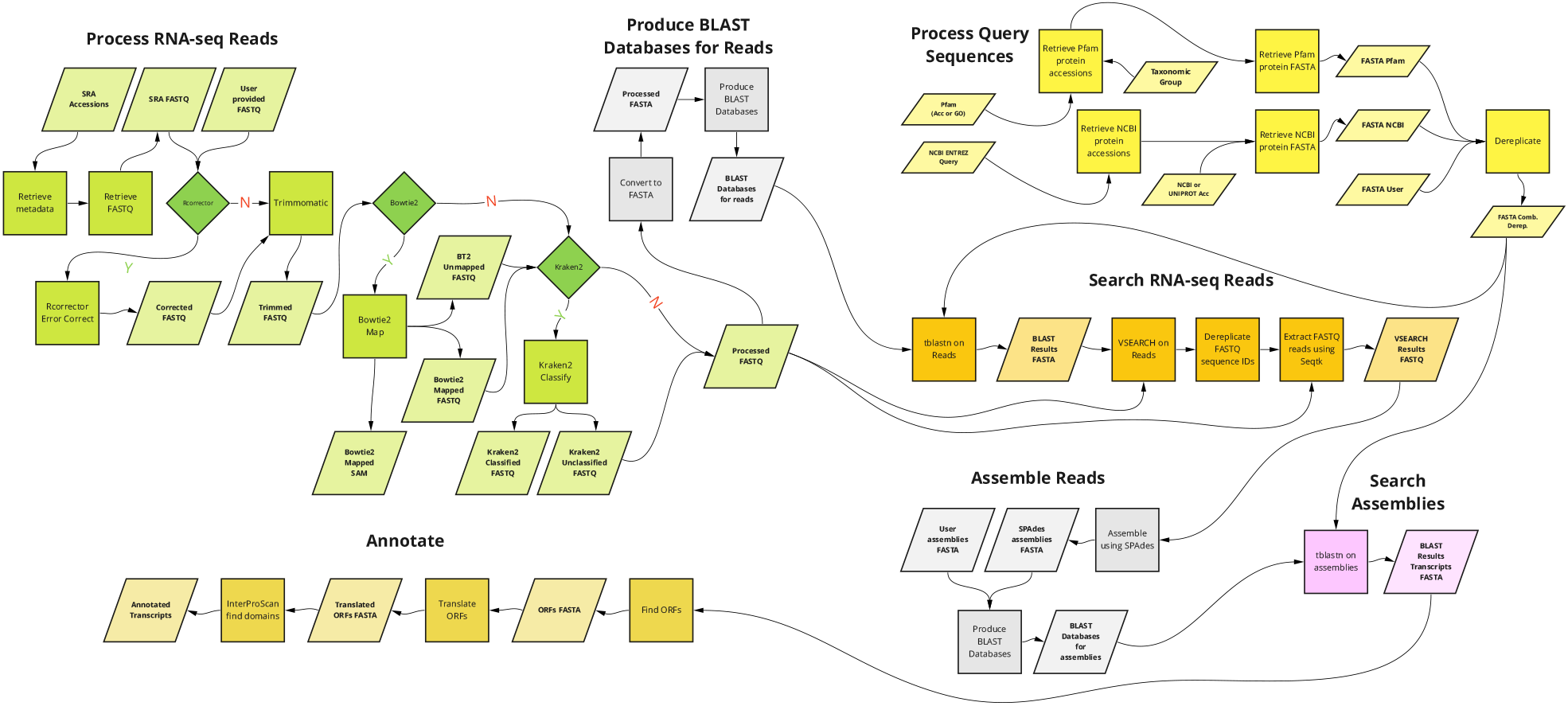
Overview of the main components of kakapo workflow.

**Figure 2:**
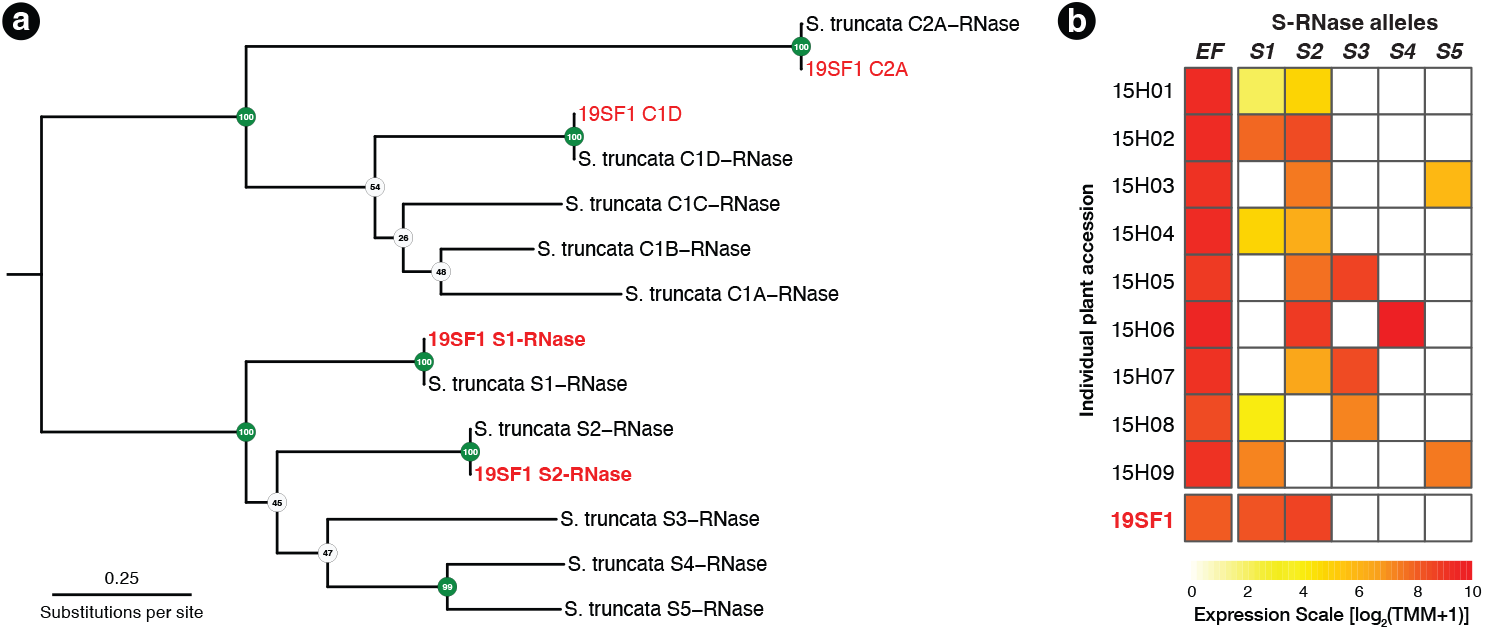
**(a)** A RAxML gene tree of *Schlumbergera truncata* T2/S-RNases from Ramanauskas and Igić (2021) and KAKAPo-recovered sequences from reads from style tissue RNA-seq of sample 19SF1. Two new S-RNases are recovered, labeled Si and S_2_. Other genes in the gene family are also found, labeled C1D and C2A, but these belong members of this large gene family with functions unrelated to self-incompatibility (non-S). **(b)** Summary of S-RNase expression data for nine *S. truncata* individuals discussed in Ramanauskas and Igić (2021), as well as 19SF1, from which two S-allele sequences are extracted by kakapo. Individual plant ID accessions are shown in rows on the left. Control gene Elongation factor 1-*α* 1 (EF) expression is presented in the first column. Pistil-expressed S-RNase (S_1_ – S_5_) follows the expected pattern. Each genotype is heterozygous (expresses two S-RNase alleles).

### Cactus Virus X in 19SF1

54,282 RNA-seq reads from individual 19SF1 mapped to the 99.5% of the 6,614 bp Cactus Virus X reference sequence NC_002815.2. One hundred or more reads mapped to 96.13% and fifty or more reads mapped to 98.53% of the reference sequence (Figure 3a). The genome we recovered is closely related to three CVX sequences (LC128411, KM365479, and AF308158) and not Zygocactus virus X, Schlumbergera virus X, or Pitaya virus X, three of several strains or species in this group (Figure 3b).

**Figure 3:**
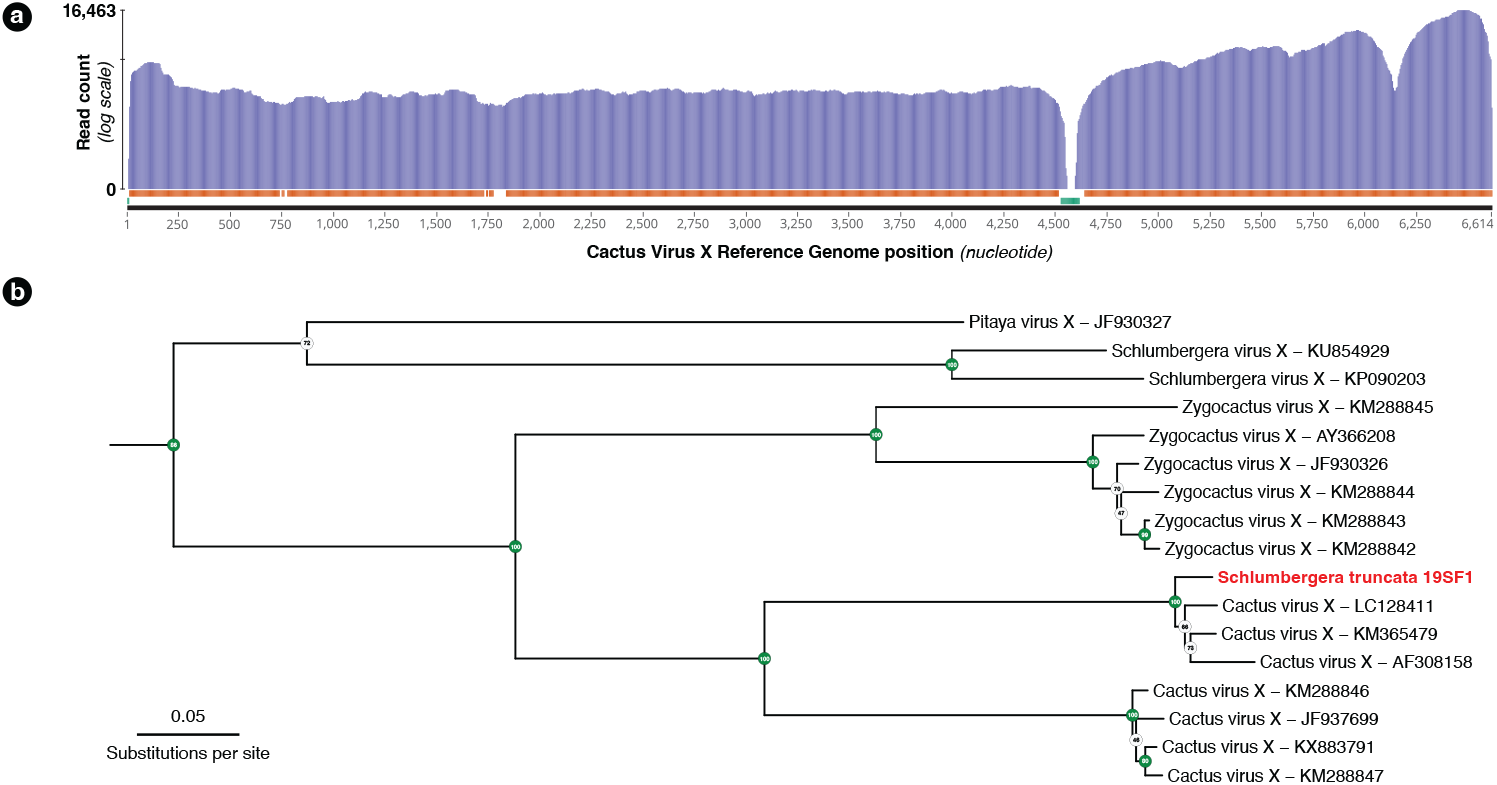
**(a)** Coverage plot of the 54,282 *Schlumbergera truncata* RNA-seq reads from sample 19SF1 mapped to Cactus Virus X (CVX) genome. Regions indicated below with the orange bar have coverage of 100 or more reads (96.13%) and those with the green bar have fewer than 50 reads (1.47%). The apparent gap just after nucleotide 4,500 likely reflect a deletion in 19SF1. **(b)** A RAxML tree of a (previously) unknown pathogen and closely related genome sequences. kakapo-recovered sequences (reads from style tissue RNA-seq of sample 19SF1) and Gen-Bank reveal that our sample contains a strain of CVX.

## Discussion

Kakapo allows users to extract and assemble a specified gene or gene family from any number of their own RNA-seq data or deposited sequence accessions. Its utility is considerable, because studies across life sciences increasingly rely on RNA-seq data, and datasets deposited to the NCBI Sequence Read Archive (SRA) are proliferating. In addition to serving as an archive for the original studies, these datasets present an opportunity for novel research, particularly in a variety of research programs concerned with the evolution of gene families. Kakapo can be deployed in a variety of fields for extraction of particular genes in studies of gene function and evolution, or (for instance) in phylogenetics and systematics to extend coverage for studies that rely on hyb-seq (Johnson et al. 2019).

The key feature of kakapo is the ease with which analyses of additional samples can be performed. The example dataset described above is small, it includes only one sample, facilitating the ease of distribution with kakapo. A user may simply add more FASTQ file paths or SRA accessions to the project configuration file, and rerun the analysis. Once the configuration files are created and tested on one sample, kakapo is intended to be used in a “set it and forget it” manner. The search strategy files are completely reusable. They are written once by the user, parameters tuned, if needed, and simply reused later. In most cases, only the project name, output directory, SRA accessions, and FASTQ file list will differ between projects. This should be remarkably convenient for repeated inquiry in a specific gene family or families (a common theme of research), and provide a straightforward way to ensure easily reproducible research. Finally, kakapo design allows easy expansion. In future work. we intend to implement support for alternative software (mappers, assemblers, etc.), to allow users to choose a tool that may be better suited for them.

## Acknowledgments

This work was supported in part by the National Science Foundation grant NSF-DEB-1655692 to B.I. We ask users of the pipeline to contribute to the Kākāpō Recovery program, aimed at helping save the eponymous critically endangered species: https://www.doc.govt.nz/our-work/kakapo-recovery/get-involved/donate.

## References

Altschul, S. F., W. Gish, W. Miller, E. W. Myers, and D. J. Lipman, 1990. Basic local alignment search tool. Journal of Molecular Biology 215:403–410.

Bateman, A., E. Birney, L. Cerruti, R. Durbin, L. Etwiller, S. R. Eddy, S. Griffiths-Jones, K. L. Howe, M. Marshall, and E. L. L. Sonnhammer, 2002. The Pfam Protein Families Database. Nucleic Acids Research 30:276–280.

Bolger, A. M., M. Lohse, and B. Usadel, 2014. Trimmomatic: a flexible trimmer for Illumina sequence data. Bioinformatics 30:2114–2120.

Eisen, J. A., D. Kaiser, and R. M. Myers, 1997. Gastrogenomic delights: A movable feast. Nature Medicine 3:1076–1078.

Handelsman, J., M. R. Rondon, S. F. Brady, J. Clardy, and R. M. Goodman, 1998. Molecular biological access to the chemistry of unknown soil microbes: a new frontier for natural products. Chemistry & biology 5:R245–R249.

Johnson, M. G., L. Pokorny, S. Dodsworth, L. R. Botigue, R. S. Cowan, A. Devault, W. L. Eiserhardt, N. Epitawalage, F. Forest, J. T. Kim, et al., 2019. A universal probe set for targeted sequencing of 353 nuclear genes from any flowering plant designed using k-medoids clustering. Systematic Biology 68:594–606.

Jones, P., D. Binns, H.-Y. Chang, M. Fraser, W. Li, C. McAnulla, H. McWilliam, J. Maslen, A. Mitchell, G. Nuka, S. Pesseat, A. F. Quinn, A. Sangrador-Vegas, M. Scheremetjew, S.-Y. Yong, R. Lopez, and S. Hunter, 2014. InterProScan 5: genome-scale protein function classification. Bioinformatics 30:1236–1240.

Katz, K., O. Shutov, R. Lapoint, M. Kimelman, J. R. Brister, and C. O’Sullivan, 2022. The Sequence Read Archive: a decade more of explosive growth. Nucleic acids research 50:D387–D390.

Kodama, Y., M. Shumway, and R. Leinonen, 2012. The Sequence Read Archive: explosive growth of sequencing data. Nucleic Acids Research 40:D54–D56.

Kreuze, J. F., A. M. Vaira, W. Menzel, T. Candresse, S. K. Zavriev, J. Hammond, K. H. Ryu, and I. R. Consortium, 2020. ICTV Virus Taxonomy Profile: Alphaflexiviridae. The Journal of General Virology 101:699–700.

Langmead, B. and S. L. Salzberg, 2012. Fast gapped-read alignment with Bowtie 2. Nature Methods 9:357–359.

Li, H., 2012. seqtk, toolkit for processing sequences in fasta/q formats. URL https://github.com/lh3/seqtk.

Quast, C., E. Pruesse, P. Yilmaz, J. Gerken, T. Schweer, P. Yarza, J. Peplies, and F. O. Glöckner, 2013. The SILVA ribosomal RNA gene database project: improved data processing and web-based tools. Nucleic Acids Research 41:D590–D596.

Ramanauskas, K. and B. Igić, 2021. RNase-based self-incompatibility in cacti. New Phytologist 231:2039–2049.

Rognes, T., T. Flouri, B. Nichols, C. Quince, and F. Mahé, 2016. VSEARCH: a versatile open source tool for metagenomics. PeerJ 4:e2584.

Sánchez-Cabrera, M., F. J. Jiménez-López, E. Narbona, M. Arista, P. L. Ortiz, F. J. Romero-Campero, K. Ramanauskas, B. Igić, A. A. Fuller, and J. B. Whittall, 2021. Changes at a critical branchpoint in the anthocyanin biosynthetic pathway underlie the blue to orange flower color transition in Lysimachia arvensis. Frontiers in Plant Science 12:633979.

Song, L. and L. Florea, 2015. Rcorrector: efficient and accurate error correction for Illumina RNA-seq reads. GigaScience 4:48.

Taylor, N. and D. Zappi, 2017. Schlumbergera truncata. The IUCN Red List of Threatened Species 2017:e.T152554A121599528.

Wood, D. E., J. Lu, and B. Langmead, 2019. Improved metagenomic analysis with Kraken 2. Genome Biology 20:257.

